# Genomic location dictates lytic promoter activity during herpes simplex virus latency

**DOI:** 10.1101/2023.12.05.570201

**Authors:** Navneet Singh, Sherin Zachariah, Aaron T. Phillips, David Tscharke

## Abstract

Herpes simplex virus 1 (HSV-1) is a significant pathogen that establishes life-long latent infections with intermittent episodes of resumed disease. In mouse models of HSV infection, persistent low-level lytic gene expression has been detected during latency in the absence of spontaneous reactivation events leading to new virus production. This viral activity during latency has been reported using a sensitive Cre-marking model for several lytic gene promoters placed in one location in the HSV-1 genome. Here we extend these findings in the same model by examining first, the activity of an ectopic lytic gene promoter in other places in the genome and second, whether native promoter activity might be detectable. We found that both for ectopic and native lytic gene promoters, *Cre* expression during latency was detected in our model, but only when the promoter was located near the ends of the unique long genome segment. This location is significant because it is in close proximity to the region from which latency associated transcripts (LAT) are derived. These results show for the first time that native HSV-1 lytic gene promoters can produce protein products during latency, but that this activity is only detectable when they are located close to the LAT locus.

**Author summary:** HSV is a significant human pathogen and the best studied model of mammalian virus latency. Traditionally the active (lytic) and inactive (latent) phases of infection were considered to be distinct, but the notion of latency being entirely quiescent is evolving due to the detection of some lytic gene expression during latency. Here we add to this literature by finding that activity can be found for native lytic gene promotors as well as for constructs placed ectopically in the HSV genome. However, this activity was only detectable when these promoters were located close by a region known to be transcriptionally active during latency. These data have implications for our understanding of HSV gene regulation during latency and the extent to which transcriptionally active regions are insulated from adjacent parts of the viral genome.

## Introduction

HSV-1 infects around 67% of world population and generally causes mild disease in the form of cold sores (Looker et al. 2015). For most individuals HSV-1 disease is infrequent and often asymptomatic, however that is not always the case and the virus can cause severe diseases such as neonatal herpes, and herpes keratitis and encephalitis. Initial HSV-1 infection usually begins in the skin or mucosal epithelium, during which expression of viral genes occurs in a sequential cascade, categorised as immediate-early (IE), early (E) and late (L), ultimately resulting in the production of new virus particles (Honess and Roizman 1974, Barklie Clements, Watson and Wilkie 1977). The late genes are further divided into two subclasses as leaky-late (γ_1_) or true-late (γ_2_), where the expression of latter strictly occurs following DNA replication (Holland et al. 1980, Conley et al. 1981). The virus quickly spreads to the cell bodies of innervating neurons within the sensory or autonomous nervous system, where latency is established allowing viral persistence. Virus replication may occur in neurons during productive infection, but it is not necessary for the establishment of latency (Speck and Simmons 1991, Speck and Simmons 1992). During latency, the HSV-1 genome persists in a non-productive state within the latently-infected neurons, while retaining the potential for reactivation.

The HSV-1 genome as it is packaged in the virion has two segments of unique sequence, termed unique long (U_L_) and unique short (U_S_), each of which is bracketed by repeats at each end (R_L_ and R_S_). By convention, the genome is considered to start with the long segment and genes are numbered from left to right in U_L_ and U_S_ (Fig 1A). To complete this nomenclature, the repeats that are at the ends of the genome are referred to as being terminal (TR_L_ and TR_S_) and those that join the long and short segments are internal (IR_L_ and IR_S_). Once inside a host cell, the genome becomes a circle and the long and short segments are able to flip with respect to each other, so there is no biological distinction between internal and terminal repeats. These repeated regions are large enough to encode some genes, meaning they are present in two copies, with each copy adjacent to different genes from U_L_ or U_S_.

**Figure 1:**
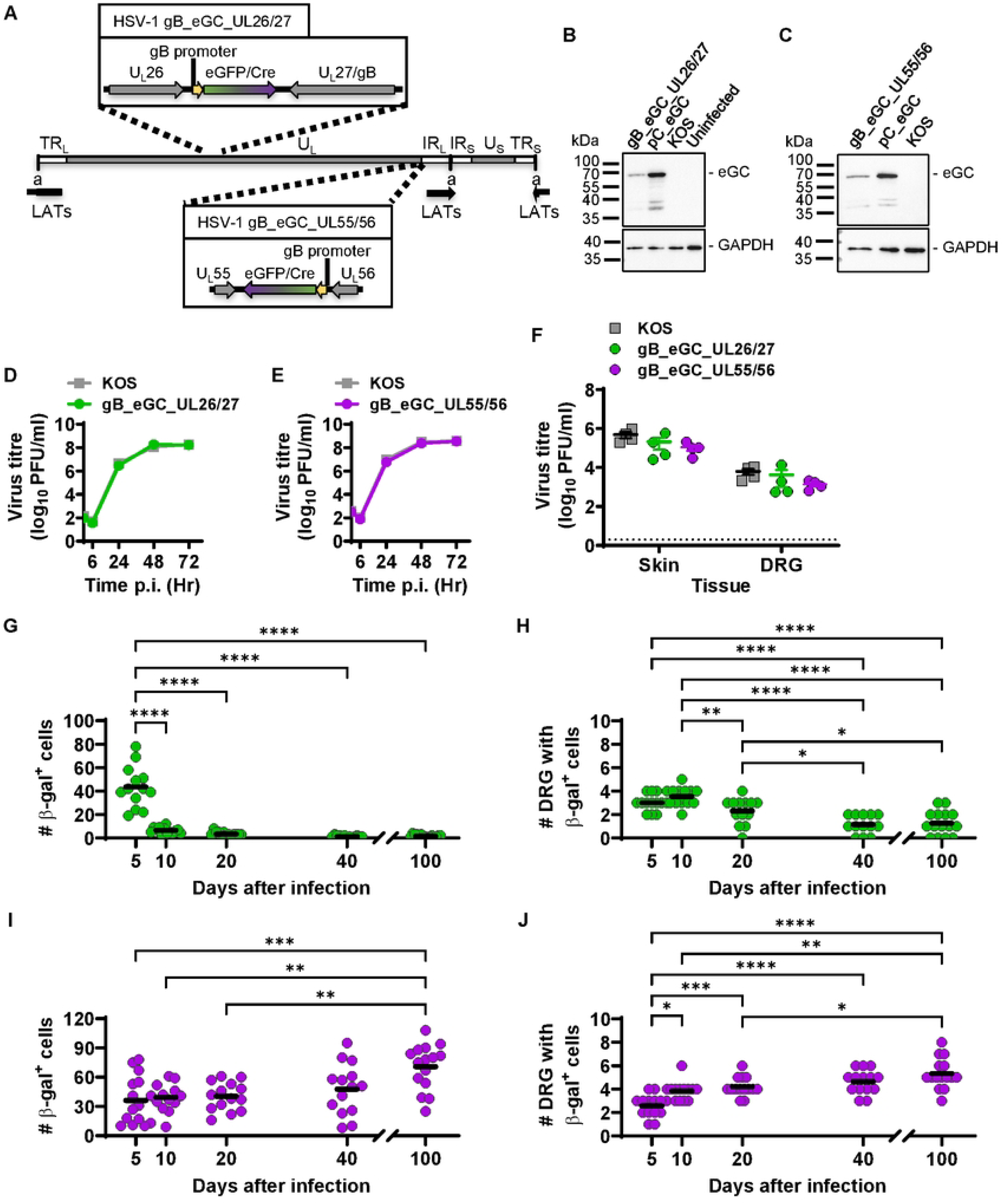
Activity of ectopic gB promoter from two independent locations. (A) Schematic showing HSV-1 genome (middle; to scale) with detail of the insertions made in HSV-1 gB_eGC_UL26/27 (top) and gB_eGC_ULSS/56 (bottom) respectively. (B,C) Detection of Cre in Vero cells infected with viruses as shown by western blotting; GAPDH was used as a loading control and sizes are shown to the left. (D.E) Replication of recombinants assessed in comparison with parent in Vero cells. Data are shown as mean ± SEM from three replicates (error bars are obscured by data points). (F) Mice were infected on the flank with recombinants and parent viruses and viral loads were measured in skin and DRG after 5 days. Each symbol represents one mouse and dotted line signifies limit of detection (2 PFU/sample). (G-J) Groups of ROSA26R mice were infected with HSV-1 gB_eGC_UL26/27 (G,H) or gB_eGC_UL55/56 (I.J), and their DRG collected and stained for J3-gal expression. Data are pooled from two independent experiments for each virus. The number of J3-gal+ cells per mouse (G,I) and the number of DRG with at least one J3-gal+ cell (H.J) are shown. Differences in means were compared using one-way ANOVA with Bonferroni’s post-test to calculate pa1rw1se comparisons(*. *P<0.05;* **, P<0.01; ***, P<0.001; unmarked, not significant).

HSV-1 latency at the level of whole organism is largely quiescent as spontaneous reactivation events are exceptionally rare (Laycock et al. 1991, Feldman et al. 2002, Gebhardt and Halford 2005). However, this perspective is challenged when viral activity is examined at the cellular level. High level gene expression during latency is limited to non-coding RNAs known as latency-associated transcripts (LATs) and certain micro-RNAs that originate from the LAT region. The expression of LATs begins early during infection, which aligns with the simultaneous establishment of latency and productive infection at the cellular level (Speck and Simmons 1992) (Spivack and Fraser 1988). Studies employing careful PCR and *in situ* hybridisation analysis have revealed that high levels of LATs are found in only 5 to 30% of latently-infected neurons at a given time in latency (Mehta et al. 1995, Maggioncalda et al. 1996, Ellison et al. 2000, Chen et al. 2002b, Wang et al. 2005a). Several reports over the years have provided evidence supporting the presence of lytic transcripts in the latently-infected trigeminal ganglia of mice, challenging the traditional view of strict viral latency (Green, Courtney and Dunkel 1981, Kramer and Coen 1995, Tal-Singer et al. 1997, Kramer et al. 1998, Chen et al. 2002a, Feldman et al. 2002, Pesola et al. 2005, Maillet et al. 2006, Ma et al. 2014). One notable report showed that lytic genes belonging to one of the classes were expressed in almost two-thirds of infected neurons (Ma et al. 2014), a far higher frequency than neurons exhibiting spontaneous reactivation (Margolis et al. 2007). This suggests that lytic gene transcription is a common phenomenon during latency in the absence of overt reactivation and is likely to be biologically relevant as an increase in viral activity were correlated with a progressive response from the host (Ma et al. 2014).

There is also evidence that some lytic transcripts may generate protein during latency, which is from two main sources. First, immune infiltrates consisting primarily of virus-specific and activated CD8^+^ T cells have been found in latently-infected sensory ganglia (Halford, Gebhardt and Carr 1996, Khanna et al. 2003, van Lint et al. 2005, van Velzen et al. 2013, Feldman et al. 2002, Sawtell 2003, Margolis et al. 2007). Second, the use of Cre reporter mice, such as ROSA26R (Soriano 1999), infected with recombinant viruses that express *Cre*-from lytic gene promoters suggests these promoters are active during latency. In the ROSA26R model, any Cre expression leads to the indelible marking of neurons and acts as a record of historic gene expression (Proenca et al. 2008, Wakim et al. 2008). In these experiments, accumulation of marked neurons was observed during latency indicating that lytic gene promoters were able to drive *Cre* expression beyond the acute infection. Promoters for infected cell protein (ICP)47 (U_S_12), ICP6 (U_L_39), and glycoprotein B (gB) (U_L_27) were able to drive Cre expression during latency, but not ICP0 (Russell and Tscharke 2016). Thus, the expression of lytic gene expression during latency could stem from unsuccessful abortive reactivation events by the virus, incomplete repression of the genome, or more speculatively, a latency-associated gene expression program (Singh and Tscharke 2020).

It is important to acknowledge a significant limitation of the study conducted by Russell and Tscharke, which is that the promoters were not examined in their natural location (Russell and Tscharke 2016). A recent study investigating the ICP47 promoter activity in latency using the same model found no evidence of Cre expression from either its native location or from the intergenic region between U_L_26 and U_L_27 (U_L_26/U_L_27). Importantly however, this study recapitulated previous data showing expression from the ICP47 promoter in latency when placed between U_L_3 and U_L_4 (U_L_3/U_L_4), using the original and several newly made viruses (Russell and Tscharke 2016, Velusamy et al. 2023). These data led to the speculation that the placement of this promoter close to the LAT promoter, which is active during latency might be required for expression to be detected during latency.

Here we have systematically investigated whether the expression of lytic genes during latency is influenced by the location of genes within HSV-1 genome, using an extended set of ectopic and native lytic gene promoters. This was accomplished by utilizing expression constructs inserted into multiple sites within the genome and then examining native promoter activity at sites found to be permissive for ectopic expression constructs. The results show the first evidence of gene expression from native HSV lytic gene promoters during latency and confirm that the level of this activity is influenced by promoter location in the HSV-1 genome.

## Results

### Ectopic gB promoter activity in HSV-1 latency depends on genomic location

Previous studies have demonstrated that the infection of ROSA26R mice with HSV-1 recombinants expressing Cre under the control of gB and ICP47 promoters from U_L_3/U_L_4, led to promoter activation and efficient marking of neurons during latency. By contrast, the ICP47 promoter could not drive Cre expression that was detectable in ROSA26R mice during latency either from its native location (at the border of R_S_ and U_S_) or from U_L_26/U_L_27. Together these results suggest a location-dependent effect on ICP47 promoter activity during latency (Russell and Tscharke 2016, Velusamy et al. 2023). Notably, U_L_3/U_L_4 is relatively close to the LAT promoter in TR_L_, which is transcriptionally active in latency. To test whether proximity to the LAT promoter might be a determinant of expression during latency for lytic gene promoters more generally, we made two new viruses: Both had an *eGFP/Cre* fusion gene under the control of the HSV-1 gB promoter (gB_eGC) but in the first, this was located in U_L_26/U_L_27 and in the second it was placed between U_L_55 and U_L_56 (U_L_55/U_L_56) (Fig 1A). These viruses were named gB_eGC_UL26/27 and gB_eGC_UL55/56, respectively. U_L_26/U_L_27 is roughly in the middle of the U_L_ segment, far from either copy of the LAT promoter, whereas U_L_55/U_L_56 is at the far end of U_L_, close to the LAT promoter in IR_L_.

The use of U_L_26/U_L_27 as an insertion site has been shown not to compromise replication or pathogenesis (Russell, Stefanovic and Tscharke 2015), but U_L_55/U_L_56 has not been evaluated. Therefore both new viruses were characterised carefully. We confirmed the insertion of gB_eGC in the correct region for these new viruses by a whole genome digest (S2A,B Fig). Then, we checked if the eGFP/Cre fusion protein was expressed in Vero cells infected with these viruses by western blotting. Unmodified HSV-1 KOS and pC_eGC (described previously in (Russell and Tscharke 2016)) were used as negative and positive controls respectively, while GAPDH was used as a loading control (Fig 1B,C). We could detect eGFP/Cre protein, but noted that there were low molecular weight fragments at lower levels that were likely cleaved or degraded products of the protein (Fig 1B,C). Next, we validated that Cre was functional when expressed by the new viruses using a Cre-reporter cell line Vero SUA, again using HSV-1 KOS and pC_eGC as controls (Rinaldi, Marshall and Preston 1999) (S3A–S3F Fig). Finally, we showed that replication of these viruses was the same as for the parent HSV-1 KOS in Vero cells (Fig 1D,E) and in mice infected by our flank tattoo model (Fig 1F) (Russell et al. 2015).

After successfully validating these viruses we used them to investigate the activity of their ectopic gB promoters. ROSA26R mice were tattoo-infected on the flank with HSV-1 gB_eGC_UL26/27 or gB_eGC_UL55/56, their dorsal root ganglia (DRG) from thoracic 5 to lumbar 1 dermatomes were isolated, stained for β-gal activity and the number of β-gal-expressing neurons were counted (S4A,B Fig). In the case of HSV-1 gB_eGC_UL26/27, we saw a decrease in the number of β-gal^+^ cells from day 5 to 10, which remained stable thereafter (Fig 1G). However, the number of DRG with β-gal^+^ cells, which represents the spread of the virus, was similar between days 5 and 10, reduced on days 20 and 40, and remained stable thereafter in latency (Fig 1H). By contrast, HSV-1 gB_eGC_UL55/56 the β-gal^+^ cell number was similar for all times up to day 40, but was increased at day 100 compared with day 20 and earlier (Fig 1I). The number of DRG with at least one β-gal^+^ cell was also increased at day 100 compared with earlier days (Fig 1J). Thus the result for HSV-1 gB_eGC_UL55/56 is reminiscent of what was seen when the same expression cassette was placed in U_L_3/U_L_4, at a similar distance to the LAT promoter, but at the other end of U_L._ (Russell and Tscharke 2016). However, there was no evidence that the gB promoter was active during latency when driving Cre from U_L_26/U_L_27, similar to what was found for the ICP47 promoter (Velusamy et al. 2023). Together, these results demonstrate that close proximity to the LAT promoter allows lytic gene activity to be detected during HSV-1 latency in the Cre/ROSA26R marking model.

### Native gB promoter does not make protein in latency

The next aim was to check if the native promoter of gB makes protein in latency. This was achieved by combining the use of a 2A peptide with HSV-1 Cre/ROSA26R system. A 2A sequence induces steric hinderance during translation such that two proteins can be made at an equimolar ratio from a single mRNA (Donnelly et al. 2001, Atkins et al. 2007, Ahier and Jarriault 2014, Daniels et al. 2014, Tulloch, Luke and Ryan 2017, Szymczak et al. 2004). We made a recombinant virus in which the stop codon at the end of U_L_27 was replaced with sequences encoding the highly efficient T2A (Liu et al. 2017) followed by Cre (called UL27-T2A-Cre) (Fig 2A).

**Figure 2:**
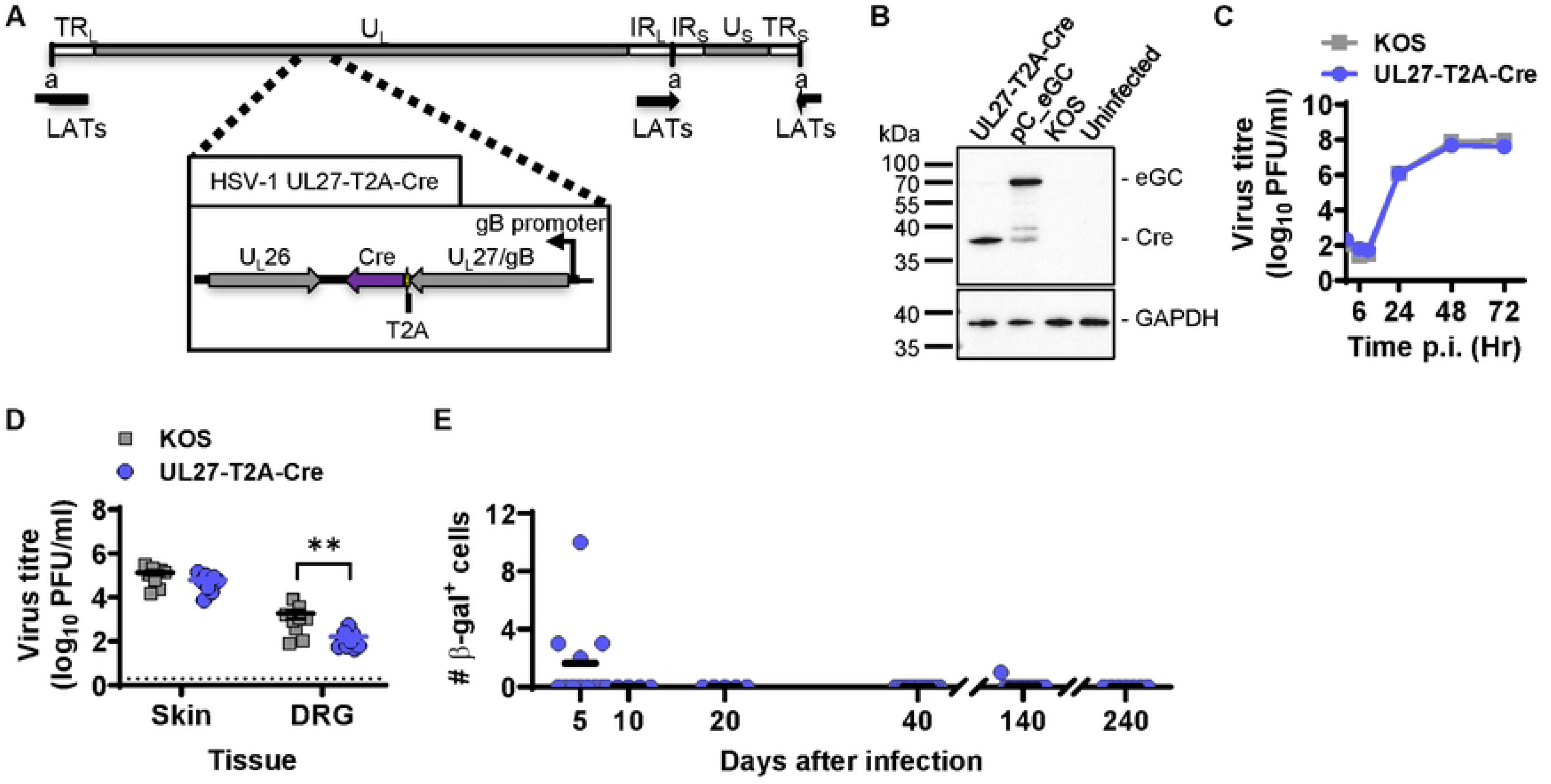
Evaluation of native gB promoter activity. (A) Structure and detail of HSV-1 UL27-T2A-Cre. (8) Detection of Cre in Vero cells infected with viruses as shown by western blotting. (C,D) Replication of HSV-1 UL27-T2A-Cre was assessed in comparison with parent *in vitro,* n=3 (C) and *in vivo* n=9 (D),. Statistical significance was determined using two-way ANOVA with Tukey’s post-test comparison(**. P<0.01; unmarked, not significant). (E) Groups of ROSA26R mice were infected with HSV-1 UL27-T2A-Cre, their DRG collected and stained for [3-gal expression. The number of [3-gal^+^ cells per mouse is shown and the data are pooled from two independent experiments. No significant differences were found between any two days.

The expression of gB and Cre as independent proteins was verified by western blotting and this virus grew at a similar rate to parent *in vitro* (Figs 2B,C). Moreover, Cre was functional in the Vero SUA reporter cells (S3 Fig). Next, we assessed the growth potential of this virus *in vivo*, in skin and DRG of C57BL/6 mice, and we found similar viral loads in the skin, however the titre of the recombinant virus was reduced in the DRG, compared with the parent virus (Fig 2D). This suggests that HSV-1 UL27-T2A-Cre may have a defect in transport to, or growth in neurons. When ROSA26 mice were infected to determine native gB promoter activity, we observed few marked neurons in four out of 11 mice on day 5, and none on days 10, 20 and 40 (Fig 2E). We decided to extend the experiment to days 140 and 240, instead of day 100 as done previously, and found that majority of the mice did not have any marked neurons on these days (Fig 2E). The lack in activity from the native gB promoter in latency is consistent with what was observed for ectopic promoter placed in the same location, but the marking at day 5 was surprisingly low. This low marking might be due to less virus in neurons, but also it is possible that the T2A sequence is not behaving as expected *in vivo*.

### T2A sequence can be used to study native promoter activity

To validate the use of T2A sequences in our HSV-1 Cre-marking model, we generated a recombinant virus with a T2A sequence between eGFP and Cre, under the control of gB promoter inserted in the U_L_3/U_L_4 region (Fig 3A). This new virus was called gB_eGTC_UL3/4 and was designed to match the previously published HSV-1 gB_eGC (Russell and Tscharke 2016), allowing a direct comparison of marking ability between these two viruses.

**Figure 3:**
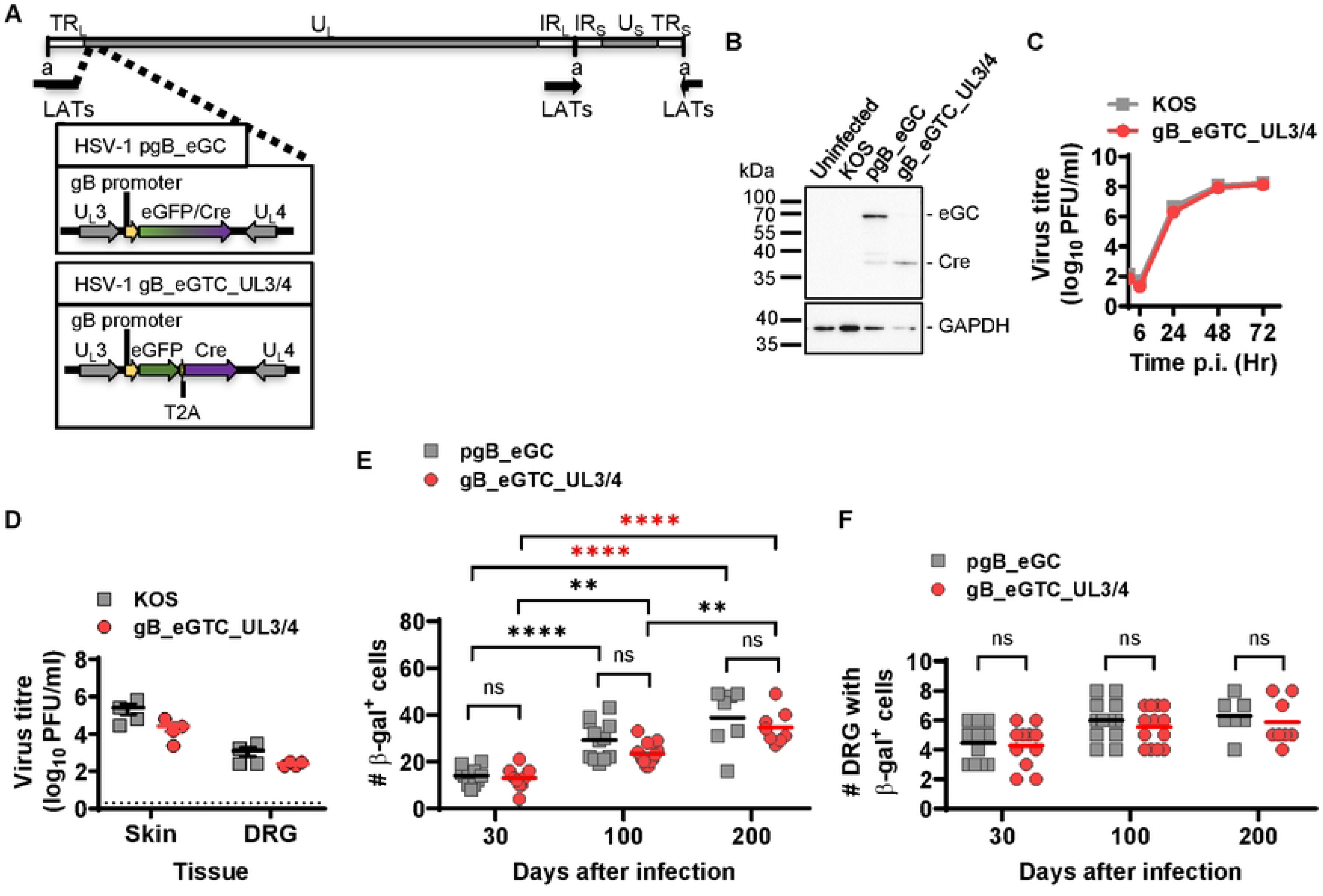
T2A does not affect neuronal marking in ROSA26R/Cre mouse model. (A) Schematic showing HSV-1 gB_eGC and HSV-1 gB_eGTC_UL3/4. (B) Detection of Cre in Vero cells infected with viruses as shown by western blotting. (C,D) Replication of HSV-1 UL27-T2A-Cre was assessed in comparison with parent *in vitro* n=3 (C), and *in vivo* n=4 (D). (E,F) Groups of ROSA26R mice were infected with HSV-1 gB_eGC or gB_eGTC_UL3/4 in parallel and their DRG collected and stained for J3-gal expression at the times shown. The number of J3-ga1^+^ cells per mouse (E) and the number of DRG with at least one J3-ga1^+^ cell (F) are shown, data were pooled from two independent experiments. Differences in means were compared using two-way ANOVA with Sidak’s post-test to calculate pairwise comparisons (**, P<0.01; ****, P<0.0001; ns, not significant; red stars signify differences of particular note).

The expression of Cre as an independent protein by the new virus was confirmed by western blotting (Fig 3B) and this virus showed similar growth kinetics to the parent both *in vitro* and *in vivo* (Figs 3C,D). To investigate the efficiency of neuronal marking when a T2A is used, groups of ROSA26 mice were infected with HSV-1 gB_eGTC_UL3/4 or HSV-1 gB_eGC. DRG were stained to count the number of β-gal-expressing neurons at days 30, 100 and 200 after infection. We found that neither the number of β-gal^+^ cells nor the number of DRG with any β-gal^+^ cells were significantly different between the viruses at any day (Fig 3E, F). Further, in accordance with previous findings there was accumulation of β-gal^+^ cells in latency regardless of presence of a T2A sequence (Fig 3E) (Russell and Tscharke 2016). Therefore, we conclude that insertion of T2A sequence does not affect neuronal marking in the ROSA26/Cre reporter system.

### Protein expression from native promoters in latency

Finally, we investigated the activity of native promoters towards the ends of U_L_ where expression of Cre from ectopic cassettes was able to be detected during latency. We chose U_L_3 and U_L_56, replacing their stop codons with a T2A, followed by Cre to make UL3-T2A-Cre and UL56-T2A-Cre, respectively (Fig 4A). Both viruses expressed Cre as an independent protein (Figs 4B,C) and had growth similar to the parent virus *in vitro* and *in vivo* (Figs 4D-G). Finally, we used these viruses to examine the activity of native promoters in the Cre marking system. When Cre was driven from the native U_L_3 promoter, we found that β-gal^+^ neurons were either non-existent or very rare, limited to one or two neurons per mouse, and to single DRG on days 5 and 10 (Fig 4H). This low level of marking was seen on days 20, 40 and 140, but there was a trend towards a higher mean and a greater fraction of mice having any marking with each later time (Fig 4H, I). At 300 days after infection, the average number of β-gal^+^ neurons and the number of DRG with marked neurons were significantly increased compared with all times up to day 40 (Figs 4H,I). Marking of neurons in ROSA26R mice by HSV-1 UL56-T2A-Cre was similar to that seen with UL3-T2A-Cre, except that an early peak was seen on day 5, before the number fell to very low levels on day 10. Thereafter there was a trend of increased means that became statistically significant compared with days 10, 20 and 40 at day 300 (Fig 4J). Similarly, number of DRG with at least one β-gal^+^ cell increased on day 300 compared with days 10 and 20 (Fig 4K). These data slow that the native U_L_3 and U_L_56 promoters can drive detectable Cre expression during HSV-1 latency.

**Figure 4:**
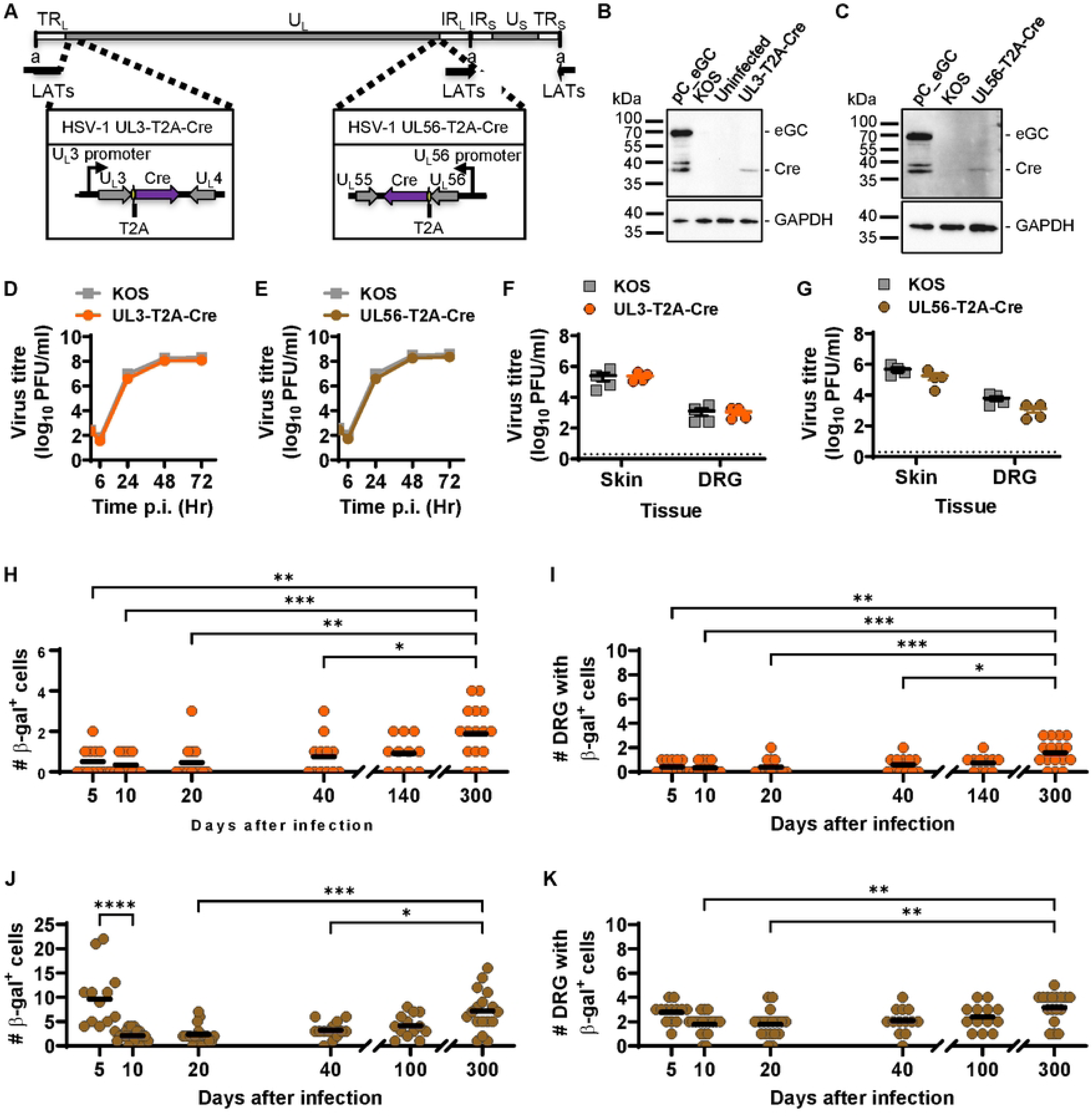
Native lytic gene promoters can express protein during HSV latency. (A) Schematic representation of HSV-1 UL3-T2A-Cre and UL56-T2A-Cre. (B,C) Detection of Cre in Vero cells infected with viruses as shown by western blotting. (D-G) Replication of recombinants was assessed in comparison with parent *in vitro,* n=3 (D,E) and *in vivo,* n=4 (F,G). (H-K) Groups of ROSA26R mice were infected with HSV-1 UL3-T2A-Cre (H,I) or UL56-T2A-Cre (J,K), their DRG collected and stained for (3-gal expression. The number of (3-gal^+^ cells per mouse (H,J) and the number of DRG with at least one (3-gal^+^ cell (l,K) are shown. Data were pooled from three independent experiments for each virus. Differences in means were compared using one-way ANOVA with Tukey’s post-test to calculate pairwise comparisons(*, *P<0.05;* **, P<0.01; ***, P<0.001; ****, P<0.0001).

## Discussion

In this study, we used a Cre/ROSA26R mouse model to investigate the influence of location of a lytic promoter in the HSV-1 genome on its activity during latency, as well as the ability of native lytic gene promoters to drive protein expression in latency. To achieve this, the gB promoter was studied in ectopic constructs from three locations: U_L_3/U_L_4, U_L_26/U_L_27 and U_L_55/U_L_56. In addition, the native promoters of gB, U_L_3 and U_L_56 were analysed for their activity by using a T2A sequence to separate the HSV protein from Cre (Karimnia et al. 2021, Nasamu et al. 2021). The results of neuronal marking by all the recombinant viruses used here have been summarised in Fig 5, showing each as percent of maximum number of neurons marked to show the differences in kinetics clearly (Fig 5). These show clearly that both for non-native and for native promoters, activity in latency was detectable from U_L_3/U_L_4 and U_L_55/U_L_56, but not from U_L_26/U_L_27.

**Figure 5:**
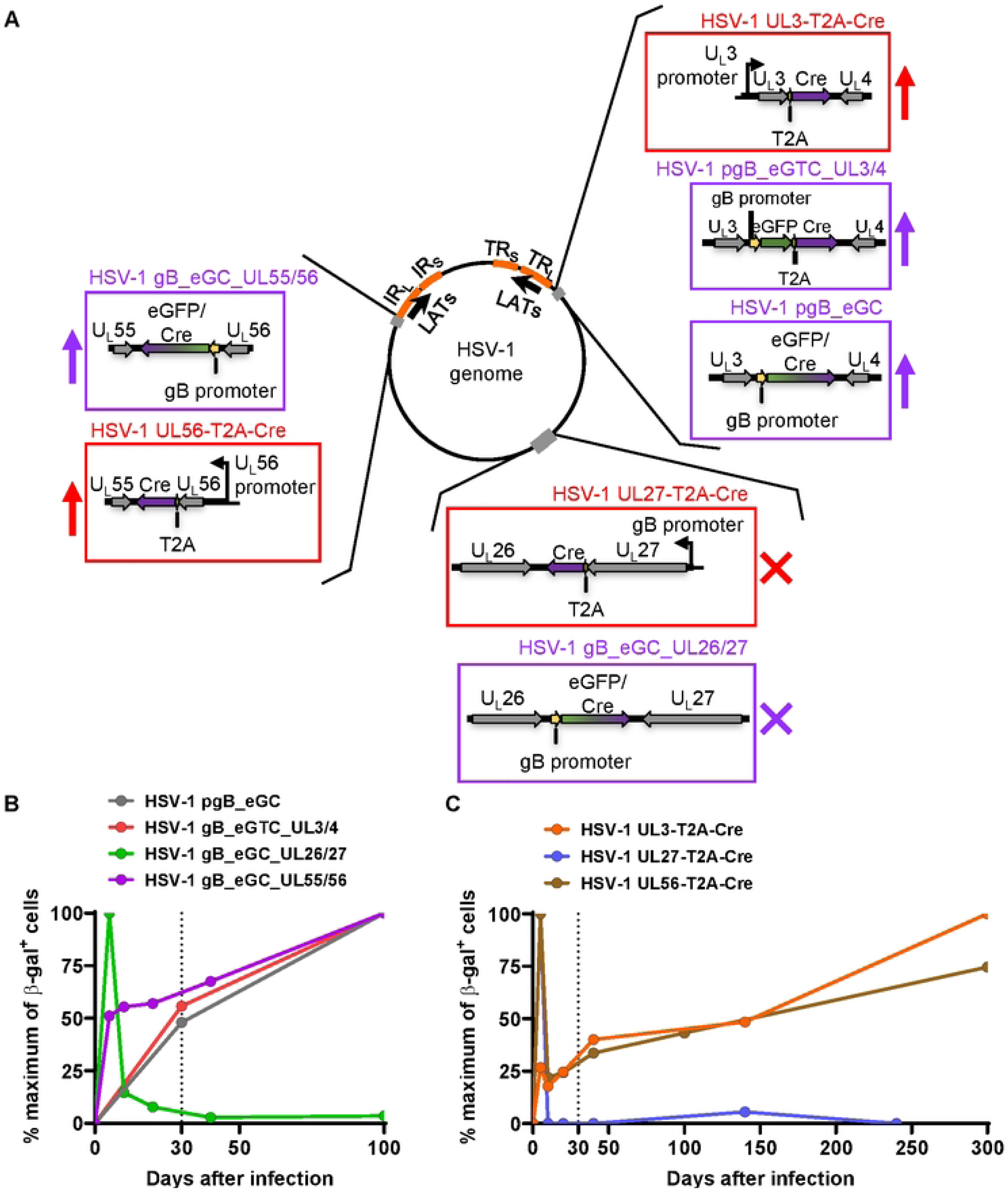
Summary of lytic promoter activity in HSV-1 latency. (A) Schematic depicting the episomal form of HSV-1 genome with the recombinants used to study ectopic (purple box) and native (red box) promoters shown. An upward arrow next to the box indicates that the promoter was active in latency whereas a cross indicates no activity was detected in latency. (B,C) The average of l3-ga1^+^ numbers in DRG of mice per time point are shown as percentage of maximum value (at the peak time point) from the viruses used to study ectopically placed gB promoter in (B) and native promoters in (C).

Prior to this study, only two promoters have been studied using ROSA26R/Cre marking models when placed in two different locations. First, there was no accumulation of β-gal+ cells in latency when Cre was driven by the ICP0 promoter, regardless of whether it was placed in the U_L_3/U_L_4 or within the U_S_5 gene (Proenca et al. 2008, Russell and Tscharke 2016). This finding suggested that while a set of other promoters was active from U_L_3/U_L_4, placement in this location alone was not adequate to guarantee activity during latency (Russell and Tscharke 2016). Second, while the promoter for ICP47 was active from U_L_3/U_L_4 during latency, this activity was not detectable from either U_L_26/U_L_27 region or U_S_12, its native location (Velusamy et al. 2023). Taken together with our new data shown here, it would seem that a location near the LAT promoter is necessary, if not sufficient to endow a lytic promoter with enough activity during latency that it can be detected in the ROSA26R/Cre model.

LATs are known to play a significant role in the maintenance of a latent state as well as reactivation (Bloom, Giordani and Kwiatkowski 2010, Phelan, Barrozo and Bloom 2017). Specifically during latency, the LAT region is characterised by the presence of euchromatic marks (Kubat et al. 2004b, Kubat et al. 2004a, Neumann et al. 2007, Wang et al. 2005b), suggesting that it may exert a permissive effect on neighbouring genes. However, considering the entire region from which RNA transcripts are regularly produced in latent neurons, it is proximity to the promoter that seems most important in leading to detectable activity during latency. The native promoter for ICP47 is within R_S_, which is within 5 kb from where the 3’ ends of the L/ST transcripts and the precursor for miR-H17 are transcribed, and yet no activity was able to be detected for this promoter during latency (Phelan et al. 2017). This is by contrast to a set of ICP47 promoter variants, all of which were active in latency when driving *eGFP/Cre* from U_L_3/U_L_4 (Velusamy et al. 2023). Considering our new data, it is interesting to note that the average number of marked neurons at day 100 was lower for gB_eGC compared with gB_eGC_UL55/56 (29 and 70, respectively). This order of marking for the gB promoter from these two locations corresponds to their distance from the LAT promoter: The gB promoter in U_L_55/U_L_56 is 2.6 kb from the start of LAT transcription whereas it is 4.1 kb for viruses that use U_L_3/U_L_4. Arguing against a simple correlation between distance from LAT and permissiveness for expression during latency, is that the fold increase in β-gal^+^ cells from the start of latency to day 100 was higher for gB_eGC in U_L_3/U_L_4 than U_L_55/U_L_56 region. This can be seen as a steeper slope for the grey compared to the purple line in Fig 5B. Some caution is required here because these comments rely on comparisons across experiments, but we note that the slope for gB_eGC is similar here as in a previous publication (Russell and Tscharke 2016) and also for the independently made gB_eGTC_UL3/4, suggesting the observations are likely to be robust. There also remains an open question as to when during infection the expression of these promoters is being affected by proximity to the LAT promoter. LATs can be expressed in some neurons from the earliest times after infection, presumably having a role in the establishment of latency and any expression at this time sets a baseline of marked neurons in the ROSA26R/Cre model.

We chose to study the gB promoter in part because activated, gB-specific T cells are found in latently-infected DRG, which strongly suggests that gB antigen is expressed during latency (Mackay et al. 2012b, Mackay et al. 2012a). However, we were not able to detect Cre expression from this promoter, either when linked to the native gB with a T2A, or from an additional copy of this promoter from the same part of the genome. Moreover, the average number of marked neurons that demonstrate survival of neurons after expression of the gB promoter at any time after infection was very low. The apparent inconsistency between the immunological studies of gB expression and ours could be attributed to the comparatively low sensitivity of our reporter system compared to the ability of T cells to detect extremely low levels of antigen. Where T cells may be able to detect as few as a handful of antigenic peptides, significantly higher levels of Cre are likely to be required to ensure migration to the nucleus, access to the target site and recombination of the *loxP* sites (Dause and Kirby 2020, Irvine et al. 2002, Purbhoo et al. 2004).

Evidence for immunological detection of lytic antigens goes beyond gB and the phenotype of resident T cells to include persistent cytokine expression by infiltrating immune cells (Chen et al. 2000). Notably, the continual cytokine expression is present in the ganglia even after infection with a thymidine kinase deficient virus that is unable to replicate and reactivate (Chen et al. 2000), suggesting that full engagement of lytic cascade is not required for the activation of immune cells. Our observation that native late gene promoters, including a true-late gene (U_L_3), can drive protein expression in latency aligns with the idea that lytic gene expression during this phase of infection is de-coupled from the ordered cascade of productive infection. Whether this activity is a component of animation phase of reactivation which never rendered complete (Kim et al. 2012, Cliffe et al. 2015, Cuddy et al. 2020, Dochnal et al. 2022), or a part of latency program of gene expression involved in maintenance of a latent state (Singh and Tscharke 2020), remains to be elucidated.

Finally, we conclude that detection of native lytic gene promoter expression during latency using ROSA26R/Cre models can be interpreted simply as showing that these promoters can be active in latent infection. It also suggests that activity is highest from promoters close to the LAT promoter and that areas of the genome at the ends of U_L_ are not entirely insulated from the de-repression of the LAT region during latency. However, the failure to detect promoter activity in latency from across the genome requires more caution in interpretation and is better considered to be falling below the limit of sensitivity of the model than to be entirely absent.

## Methods

### Cell lines and viruses

Vero cells were obtained from American Type Culture Collection (ATCC, CCL-81). Cre reporter assays were performed in Vero SUA cells (gift from Prof Stacey Efstathiou, (Rinaldi et al. 1999)). Both cell lines were cultured in minimum essential medium (MEM; ThermoFisher Scientific) supplemented with 2 or 10% heat-inactivated fetal bovine serum (FBS; Sigma-Aldrich), 4 mM L-glutamine (ThermoFisher Scientific), 5 mM HEPES buffer (ThermoFisher Scientific) and 55 µM β-mercaptoethanol (ThermoFisher Scientific). The transfections were carried out in 293A cells (ATCC, CCL-81) using Lipofectamine 2000 (ThermoFisher Scientific). 293A cells were cultured in Dulbecco’s modified Eagle medium (DMEM; ThermoFisher Scientific) supplemented with 10% heat-inactivated FBS (Sigma-Aldrich) and 2 mM L-glutamine (ThermoFisher Scientific).

HSV-1 KOS (Smith 1964) was a gift from Dr. Francis R. Carbone (University of Melbourne, Australia) and all recombinant viruses used in this study were derived from HSV-1 strain KOS. HSV-1 pC_eGC, pCmC and gB_eGC have been described previously (Russell et al. 2015) (Russell and Tscharke 2016). All viruses were titrated and grown on Vero cells as described elsewhere (Russell et al. 2015).

### Plasmid construction

The plasmid constructs used as repair template for recombinant virus generation were generated using InFusion cloning (TaKaRa). The sequence references below are based on the HSV-1 KOS genome accession number JQ673480 (Macdonald et al 2012). To construct pgB_eGC_UL26/27, promoter gB (55985 – 56282), eGFP/Cre and bovine growth hormone (BGH) polyA termination sequence were amplified from pT gB_eGC (Russell and Tscharke 2016), inserted into the *Spe*I site of pU26/7 (Russell et al. 2015). To construct pgB_eGC_UL55/56, flanking arms with U_L_55 (115549 – 116066) and U_L_56 (116067 – 116573) were amplified from HSV-1 KOS, and promoter gB, eGFP/Cre and bovine growth hormone (BGH) polyA termination sequence were amplified from pT gB_eGC (Russell and Tscharke 2016), and inserted into the *Bam*HI site of pCR bluntII vector (Invitrogen, Life Technologies) and this plasmid was used to generate HSV-1 gB_eGC_UL55/56 (S1A Fig).

To construct pgB_eGTC_UL3/4, the flanking U_L_3 and U_L_4 arms, gB promoter, eGFP and Cre-BGH polyA sequences were amplified from pT gB_eGC (Russell and Tscharke 2016). T2A sequence was synthesised as two complementary dioxynucleotides (Liu et al. 2017). These fragments were cloned into the *Bam*HI site of pCR bluntII vector (Invitrogen, Life Technologies) maintaining the original configuration as pT gB_eGC (Russell and Tscharke 2016) except that T2A was inserted in frame between eGFP and Cre.

To construct pUL27-T2A-Cre, U_L_26 (52391 – 53024) and U_L_27 (53025 – 53391) containing flanking arms were amplified from HSV-1 KOS, T2A was synthesised as two complementary dioxynucleotides (Liu et al. 2017), and Cre sequence was amplified from pT gB_eGC (Russell and Tscharke 2016). These fragments were inserted into the *Bam*HI site of pCR bluntII vector (Invitrogen, Life Technologies) and this plasmid was used to generate HSV-1 UL27-T2A-Cre (S1B Fig). To construct pUL3-T2A-Cre, T2A and Cre sequences were obtained as earlier, U_L_3 (11184 – 11610) and U_L_4 (11611 – 12049) containing fragments were amplified from HSV-1 KOS, and inserted into the *Bam*HI site of pCR bluntII vector and this plasmid was used to generate HSV-1 UL3-T2A-Cre (S1C Fig). Plasmid UL56-T2A-Cre was constructed using U_L_55 (115644 – 116144) and U_L_56 (116145 – 116660) fragments amplified from HSV-1 KOS, and T2A-Cre fragment from pUL27-T2A-Cre, all cloned into *Bam*HI site of pCR bluntII vector and this plasmid was used to generate HSV-1 UL56-T2A-Cre (S1D Fig).

The pX330 plasmid has been described previously (Cong et al. 2013) and was purchased from Addgene (plasmid 42230). The sequences coding for appropriate guide RNA were synthesised as two complementary dioxynucleotides, annealed to generate double stranded DNA fragments and inserted into the *Bbs*I site of pX330.

Oligonucleotides used to generate pX330-UL55-56 are CACCGCCAGGCGTGGTGTGAGTTTG and AAACCAAACTCACACCACGCCTGGC, pXUL26/27 are CACCTTTGTCACGGGAAAGGAAAG and AAACCTTTCCTTTCCCGTGACAAA, pX330-UL27 are CACCGCCGACGAGGACGACCTGTGA and AAACTCACAGGTCGTCCTCGTCGGC, and pX330-UL56 are CACCGACAGGGGCGCTTACCGCCAC and AAACGTGGCGGTAAGCGCCCCTGTC.

### Recombinant virus generation and *in vitro* growth analysis

All the recombinant viruses were constructed using a transfection/infection method described previously (Velusamy, Gowripalan and Tscharke 2020, Russell et al. 2015). To construct HSV-1 gB_eGC_UL26/27 or gB_eGC_UL55/56, linearised pgB_eGC_UL26/27 was cotransfected with pXUL26/27 or pgB_eGC_UL55/56 was cotransfected with pXUL26/27 respectively, into Vero cells followed by infection with HSV-1 KOS. HSV-1 gB_eGTC_UL3/4 or UL3-T2A-Cre was constructed after cotransfecting pgB_eGTC_UL3/4 or UL3-T2A-Cre respectively with pX330-mC followed by infection with HSV-1 pCmC (Russell et al. 2015). To construct HSV-1 UL27-T2A-Cre or UL56-T2A-Cre, Vero cells were cotransfected with pUL27-T2A-Cre and pX330-UL27 or pUL56-T2A-Cre and pX330-UL56 respectively, following infection with HSV-1 KOS. All the viruses used in this study have been listed in Table 1.

**Table 1.** Detail of the viruses used in this study

| Virus | Cre expression | Parent virus | Reference |
| --- | --- | --- | --- |
| HSV-1 KOS | Nil | - | (Smith 1964) |
| HSV-1 pgB_eGC | Promoter: gB<br>Location: UL3/UL4<br>intergenic region | HSV-1 KOS | (Russell and Tscharke 2016) |
| **HSV-1 pC_eGC** | Promoter: CMV IE<br>Location: UL3/UL4<br>intergenic region | HSV-1 KOS | (Russell et al.<br>2015) |
| **HSV-1 gB_eGC_UL26/27** | Promoter: gB<br>Location: U<sub>L</sub>26/U<sub>L</sub>27<br>intergenic region | HSV-1 KOS | This study |
| **HSV-1 gB_eGC_UL55/56** | Promoter: gB<br>Location: U<sub>L</sub>55/U<sub>L</sub>56<br>intergenic region | HSV-1 KOS | This study |
| **HSV-1 UL27-T2A-Cre** | Promoter: gB<br>Location: Native | HSV-1 KOS | This study |
| **HSV-1 gB_eGTC_UL3/4** | Promoter: gB<br>Location: U<sub>L</sub>3/U<sub>L</sub>4<br>intergenic region | HSV-1 pCmC<br>(Russell et al. 2015) | This study |
| **HSV-1 UL3-T2A-Cre** | Promoter: U<sub>L</sub>3<br>Location: Native | HSV-1 pCmC<br>(Russell et al. 2015) | This study |
| **HSV-1 UL56-T2A-Cre** | Promoter: U<sub>L</sub>56<br>Location: Native | HSV-1 KOS | This study |

The screening of desired recombinant was carried out based on fluorescence where possible or PCR. Each virus was purified further via three rounds of plaque purification and the absence of parent virus in the final round was confirmed by PCR. The introduced modification in the recombinant was verified by PCR, Sanger sequencing and restriction fragment length polymorphism (S2 Fig). The expression of Cre was verified by western blotting and the function was validated by Vero SUA reporter assays (S3 Fig).

The growth kinetics of newly generated viruses was checked in comparison with the parent virus KOS in Vero cells as described previously (Russell and Tscharke 2016).

### Ethics statement

All mice were sourced from Australian Phenomics Facility (APF), Canberra, Australia. This study was conducted according to ethical requirements and approval from Australian National University Animal Experimentation Ethics Committee under protocols A2017/39 and A2020/42. The research was undertaken in accordance with Australian Capital Territory Animal Welfare Act 1992 and Australian code for the care and use of animals for scientific purposes, 8th edition 2013.

### Mice and infections

At least eight weeks old, female, specific pathogen free C57BL/6 or B6.129S4-Gt(ROSA)26Sortm1So/J (ROSA26R) (Soriano 1999) were used for experiments. The mice were housed and bred at APF. ROSA26R mice were a gift from Dr. Francis R. Carbone (University of Melbourne, Australia). Mice were anaesthetised by intraperitoneal injection with Avertin (2,2,2-tribromoethanol in 2-methyl-butanol) or Ketamine/Xylazine mix before infection with 1 × 10^8^ PFU/ml of virus on the shaved flank. The procedure was followed as described previously except that the needle was charged for 20 s in the virus suspension and tattooed for a period of 20 s (Russell et al. 2015). The virus dose and infection route were same for all the experiments.

### Virus titration from skin and DRG

Mice were euthanised with a rising concentration of CO_2_, the infected area of the skin (2.5 cm vertically × 0.8 cm horizontally) and the DRG (T5-L1) on the ipsilateral side were excised, and collected in 500 µl MEM (without serum) 5 days after infection. The tissue samples were snap frozen, 5 mm diameter stainless steel bead (Qiagen) was added to all the tubes and the tissue was homogenised in TissueLyser II (Qiagen) at an oscillation frequency of 30 Hz for 90 s twice. Homogenates were subjected to three freeze/thaw cycles and the amount of infectious virus was quantified by a standard plaque assay on Vero cells.

### Detection of β-gal expression

To check expression of β-gal *in vitro*, confluent Vero SUA cells were left untreated or infected with appropriate virus at an MOI of 0.05 PFU/cell and incubated at 37 °C with 5% CO_2_ for 1 hr. The unabsorbed virus was removed, replaced with fresh medium (MEM with 2% serum) and cells were further incubated for 36 hrs. Following incubation, the cells were washed with PBS (Sigma-Aldrich), fixed with 2% paraformaldehyde (Electron Microscopy Sciences)/ 0.5% glutaraldehyde (Sigma-Aldrich) (in PBS) for 4 hrs at 4 °C, washed with PBS again and incubated in the permeabilization solution (2 mM magnesium chloride (Ajax FineChem), 0.01% (w/v) sodium deoxycholate (Sigma-Aldrich), 0.02% (v/v) IGEPAL CA-630 (Sigma-Aldrich), 5 mM potassium ferrocyanide (Sigma-Aldrich) and 5 mM potassium ferricyanide (Sigma-Aldrich) in PBS) containing 1 mg/ml X-gal (Sigma-Aldrich; prepared fresh as a 40 mg/ml stock in *N,N*-Dimethylformamide (Sigma-Aldrich)) overnight at 4 °C. Cells were washed with PBS, overlayed with 50% glycerol (Sigma-Aldrich; in PBS), and then visualised and imaged using Olympus CKX41 microscope fitted with Olympus DP22 digital camera.

To enumerate β-gal-expressing cells *in vivo*, mice were euthanised with a rising concentration of CO_2_, and the DRG (T5-L1) on the ipsilateral side were collected individually in fixative as above. The DRG were incubated on ice for 1 hr and then washed twice with PBS before adding permeabilization buffer as before. Following further incubation at 4 °C for 30 mins, the solution was replaced with permeabilization buffer containing 1 mg/ml X-gal (prepared as above) and DRG were incubated at 4 °C for 12-16 hrs. DRG were washed with PBS again and incubated in 50% glycerol (in PBS) overnight. The DRG were visualised and imaged using Olympus CKX41 microscope equipped with Olympus DP22 digital camera. The β-gal^+^ cells were either counted manually or with the aid of ImageJ software (Schneider, Rasband and Eliceiri 2012).

### Restriction fragment length polymorphism

For extracting viral DNA, 80% confluent Vero cells were infected with appropriate virus in a 175 cm^2^ area flask at an MOI of 0.05 PFU/cell for 1 hr at 37 °C with 5% CO_2_. Fresh medium was added, and cells were further incubated for 48 hrs. The cells and supernatant were centrifuged to remove cell debris. Supernatant was further centrifuged at 17,684× *g* for 90 mins at 4 °C and the pellet obtained was resuspended in TE-SDS (10 mM Trizma base (pH 8.0) (Sigma-Aldrich), 1 mM EDTA (ThermoFisher Scientific) and 0.5% (w/v) SDS (Sigma-Aldrich) in water). DNA was extracted from the lysate using a standard phenol and chloroform extraction method (Sambrook and Russell 2006), digested with an appropriate restriction enzyme and electrophoresed on an agarose gel (S2 Fig).

### Western blotting

Confluent Vero cells were infected with appropriate virus at an MOI of 10 PFU/cell and incubated for 20 hrs at 37 °C with 5% CO_2_. Cells were washed once with PBS and resuspended in RIPA buffer (200 mM Trizma base (Sigma-Aldrich), 150 mM Sodium Chloride (Bacto), 1% Triton X-100 (Sigma-Aldrich), 0.5% Sodium deoxycholate (Sigma-Aldrich), 0.1% SDS (Sigma-Aldrich) and 1 Protease inhibitor tablet (Merck) per 10 ml of water, pH adjusted to 7.4). The amount of protein in the lysate was quantified using Pierce^TM^ BCA Protein Assay Kit (ThermoFisher Scientific) according to manufacturer’s instructions. 20 µg of protein was separated by sodium dodecyl sulfate-polyacrylamide gel electrophoresis (SDS-PAGE) and transferred onto Immobilon®-P PVDF membrane (Merck), followed by blocking in milk and further overnight probing at 4 °C with a primary antibody specific for Cre (1:500; Cell Signaling Technology, mAb #12830) or GAPDH (1:5000; Cell Signaling Technology, mAb #5174). A goat anti-rabbit horseradish peroxidase (HRP) conjugated secondary antibody (1:10,000; Jackson ImmunoResearch Laboratories Inc., AB_2313567) was applied to the membrane for 1 hr at room temperature. The membranes were imaged using a chemiluminescence detection system (ChemiDoc MP, Bio-Rad) after treating with a HRP substrate (Clarity Western ECL substrate, Bio-Rad) according to manufacturer’s instructions.

### Statistical analysis

Statistical comparisons of means were done using a one-way or two-way analysis of variance (ANOVA) with an appropriate post-hoc test in GraphPad Prism (Version 10.0.1). A *p* value less than 0.05 was considered significant.

## Acknowledgements

We thank Dr. Francis R. Carbone (University of Melbourne, Australia) for the gift of HSV-1 KOS and ROSA26R mice, Jon Yewdell and Jack Bennink for 293A cells and Stacey Efstathiou (NIBSC, UK) for Vero SUA cells. We thank Australian Phenomics Facility (APF) for husbandry and taking care of mice and Biomolecular Resource Facility (BRF) at the John Curtin School of Medical Research, ANU.

